## Supplementary Information for "Chromatin organization drives the exploration strategy of nuclear factors"

[†] Equally contributing authors.

**Content:**

- 7 Supplementary figures.
- 2 Supplementary movie legends.
- 2 Supplementary notes.
- 1 Supplementary table.

### Supplemental Figures

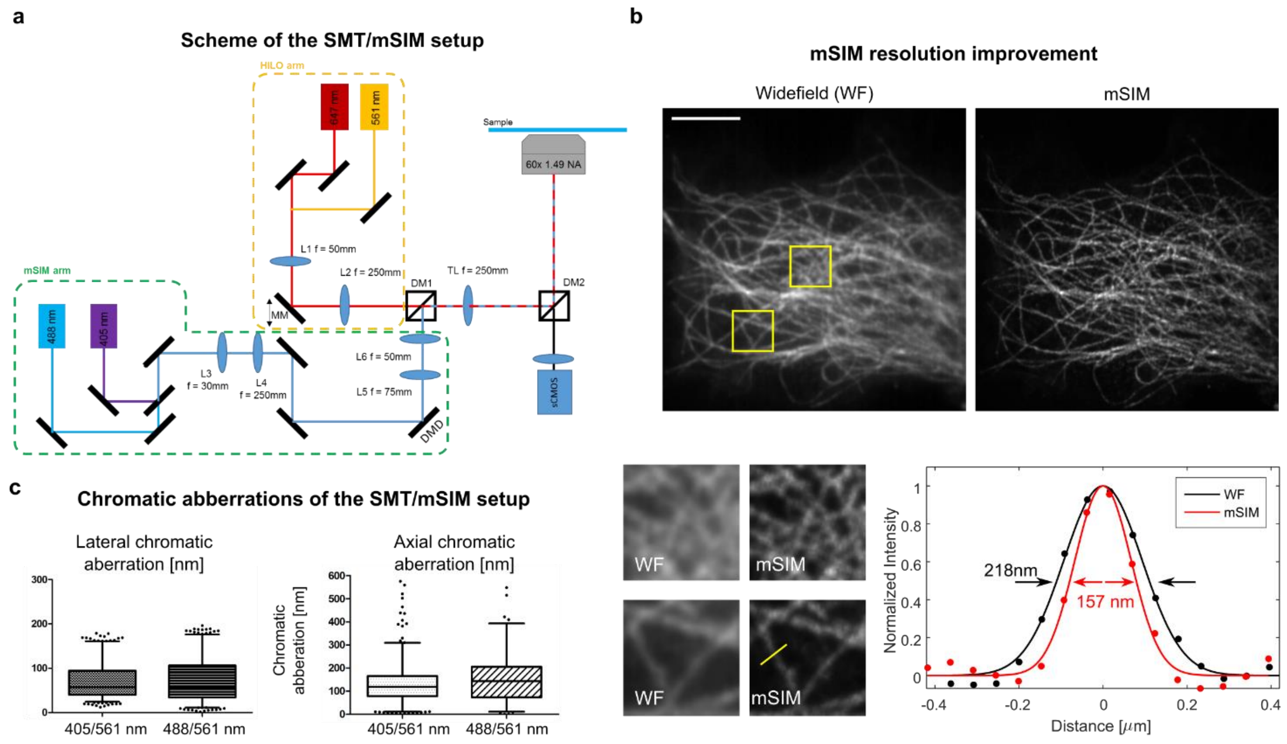


**Supplementary Figure 1. Overview of the SMT/mSIM microscope.** (**a**) Scheme of the optical setup. Two illumination arms allow for HILO and mSIM imaging of the same sample. The HILO illumination arm combines a 561nm and a 647 nm laser through a dichroic mirror, and projects a focused spot on a movable mirror (MM) that sits in a conjugated plane of the back focal plane of the microscope objective. Moving the movable mirror results in tilting the collimated beam in the front focal plane of the microscope objective. The mSIM illumination arm combines a 405 nm and a 488 nm laser through a dichroic mirror. The beam is then expanded by the couple of lenses L3 and L4 and directed to a digital micromirror device (DMD) that allows to project arbitrary patterns on the sample plane. The two illumination arms are combined through a dichroic mirror (DM1, Di03-R488-t3-25x36, Semrock Inc), and projected on the back aperture of the objective by an Olympus tube lens (TL). Emitted light by the sample is separated from excitation light by the dichroic mirror DM2 (Di03-R405/488/561/635-t3-25x36, Semrock inc.). The collected light is projected on a sCMOS camera (Orca Fusion, Hamamatsu Photonics). (**b**) The increase in lateral resolution of the mSIM microscope can be appreciated by imaging microtubules in fixed cells. Shown is the line profile along the segment highlighted in yellow. (**c**) We characterized chromatic aberrations between the HILO channel (561nm) and the mSIM channels (405 or 488 nm) by imaging 100nm microspheres (Tetraspeck Beads, ThermoFisher). Both lateral and axial chromatic aberrations were found to be below the resolution limit of mSIM.

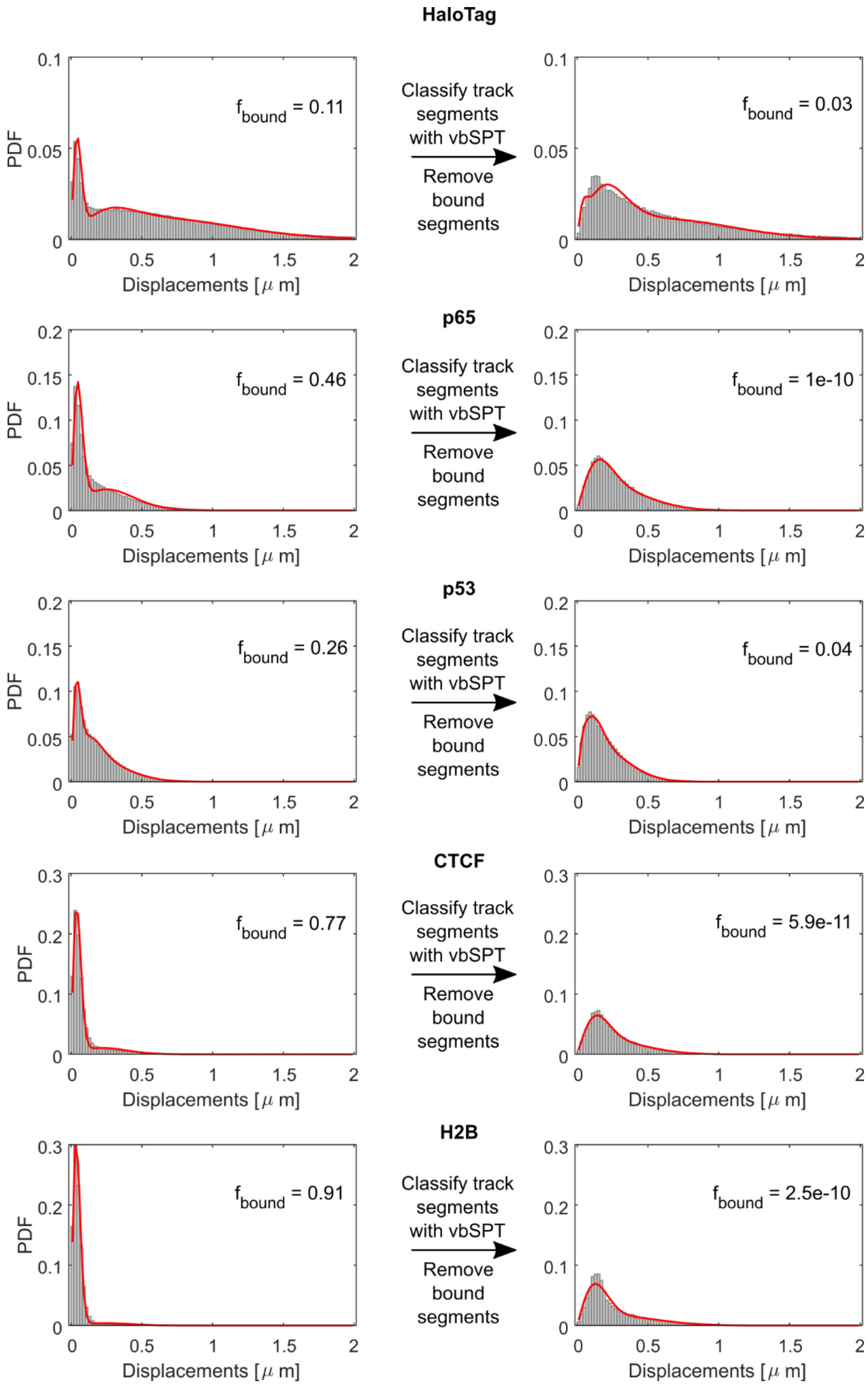


**Supplementary Figure 2. Filtering out chromatin bound molecules by vbSPT.** We use vbSPT to classify track segments into bound and diffusing components, and then filtered out the segments identified as bound molecules, in order to focus the analysis of diffusional anisotropy on the diffusing components only. To check that vbSPT successfully identified bound segments, we reanalyzed the distribution of displacements for each of the factors, after discarding those bound segments. In every case the residual bound fraction was estimated to be less than 5%. Source data: same as Figure 2.

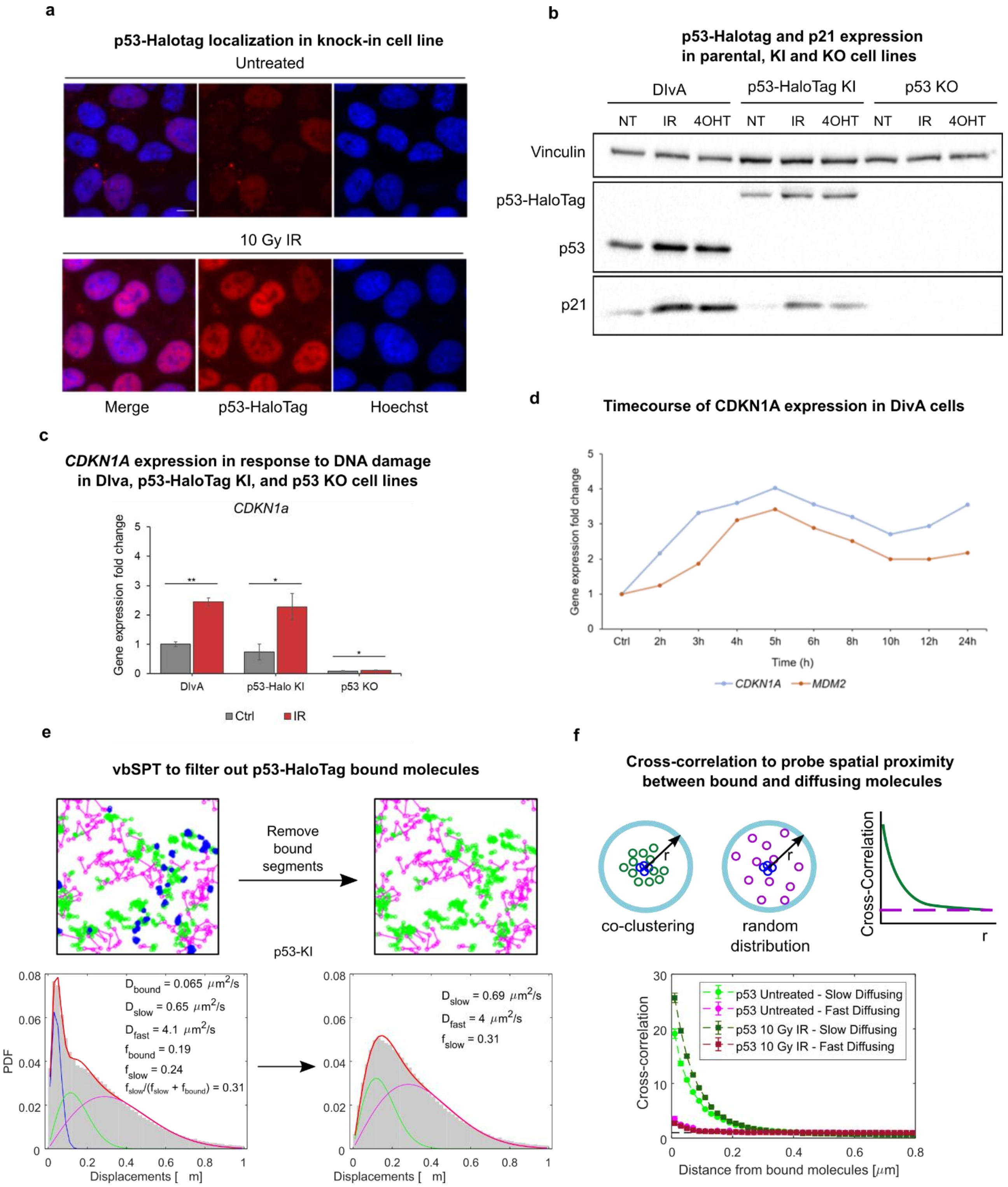


**Supplementary Figure 3. Characterization of the p53-HaloTag knock-in (KI) cell line.** (**a**) HaloTag-p53 displays the expected nuclear localization and accumulation in response to activation by 10Gy IR. (**b**) Western-blot analysis reveals that the p53-HaloTag KI cell line displays accumulation of p53 and its target p21 upon activation by 10Gy IR or Tamoxifen, while the p53-knock out (KO) cell line does not. (**c**) Analysis of p53 target gene expression by RT-PCR displayed similar induction of p53 target genes in parental cells and p53-Halotag KI cells and no induction in p53-KO cells (error bars: SD, $N_{replicates}$ = 3, statistical test: Student's t-test). (**d**) Time-course of target gene expression for *CDKN1A* and *MDM2* in p53-Halotag KI highlights a peak in p53 transcriptional activity between 4 and 5 hours post irradiation. (**e**) vbSPT can be used to correctly filter out the bound population of p53 molecules. (**F**) Co-clustering of p53 bound and diffusing molecules by cross-correlation analysis. Slow diffusing p53 molecules tend to co-cluster more frequently with bound molecules than fast ones (Data same as in Figure 3).

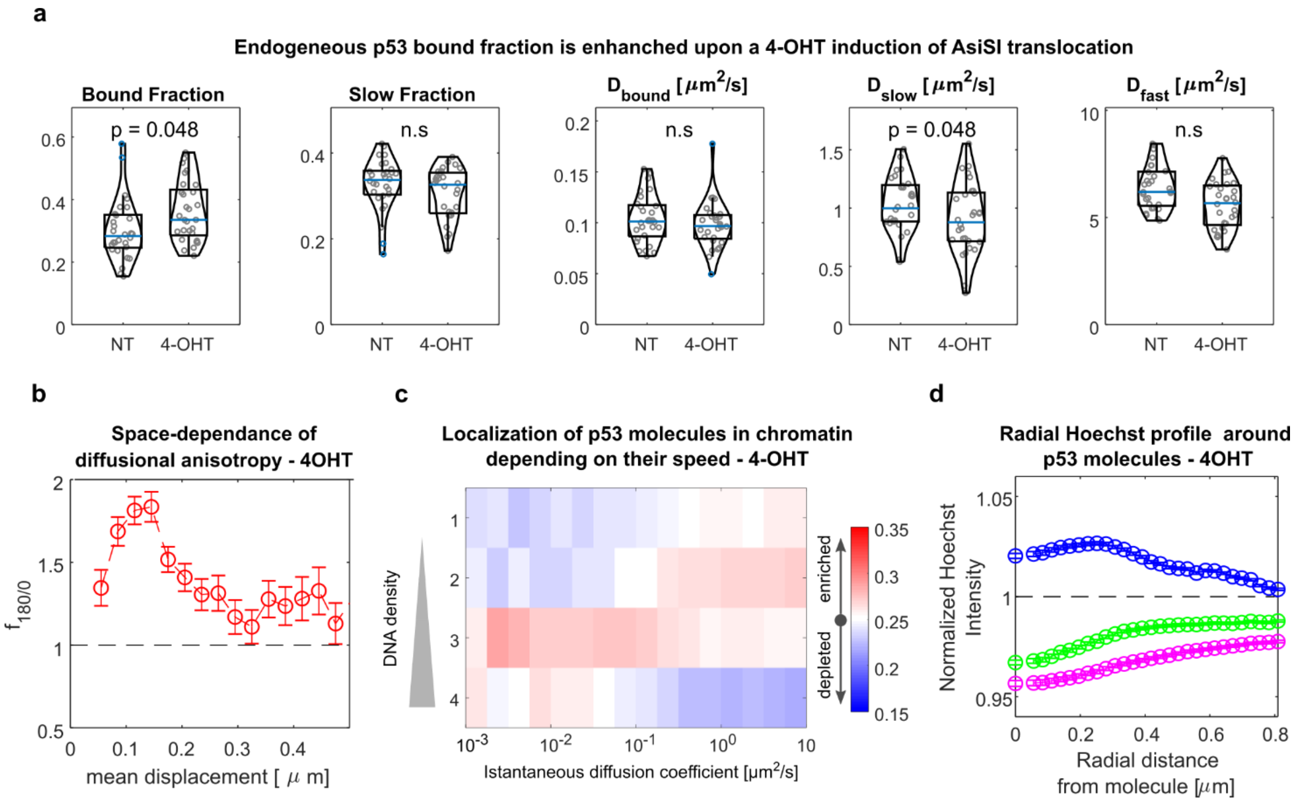


**Supplementary Figure 4. Analysis of p53-HaloTag diffusion upon induction of DNA damage by AsiSI activation in DIvA p53-HaloTag knock-in cells.** DIvA cells were treated with Tamoxifen (4-OHT) for 4 hrs to induce AsiSI translocation in the cell nucleus and analyzed with our SMT/mSIM pipeline. Induction of DNA damage by AsiSI results in increased p53 bound fraction (**a**) and a diffusional anisotropy profile compatible with guided exploration (**b**). Slow diffusing/bound p53 molecules localize in regions at higher DNA density than fast diffusing molecules, as evidenced by plotting p53 localization frequency in chromatin depending on their speed **(c)** and the radial Hoechst profile around p53 molecules (**d**). ($n_{cells}$ = 29, 29 for untreated and 4-OHT respectively, statistical test Kolmogorov-Smirnov).

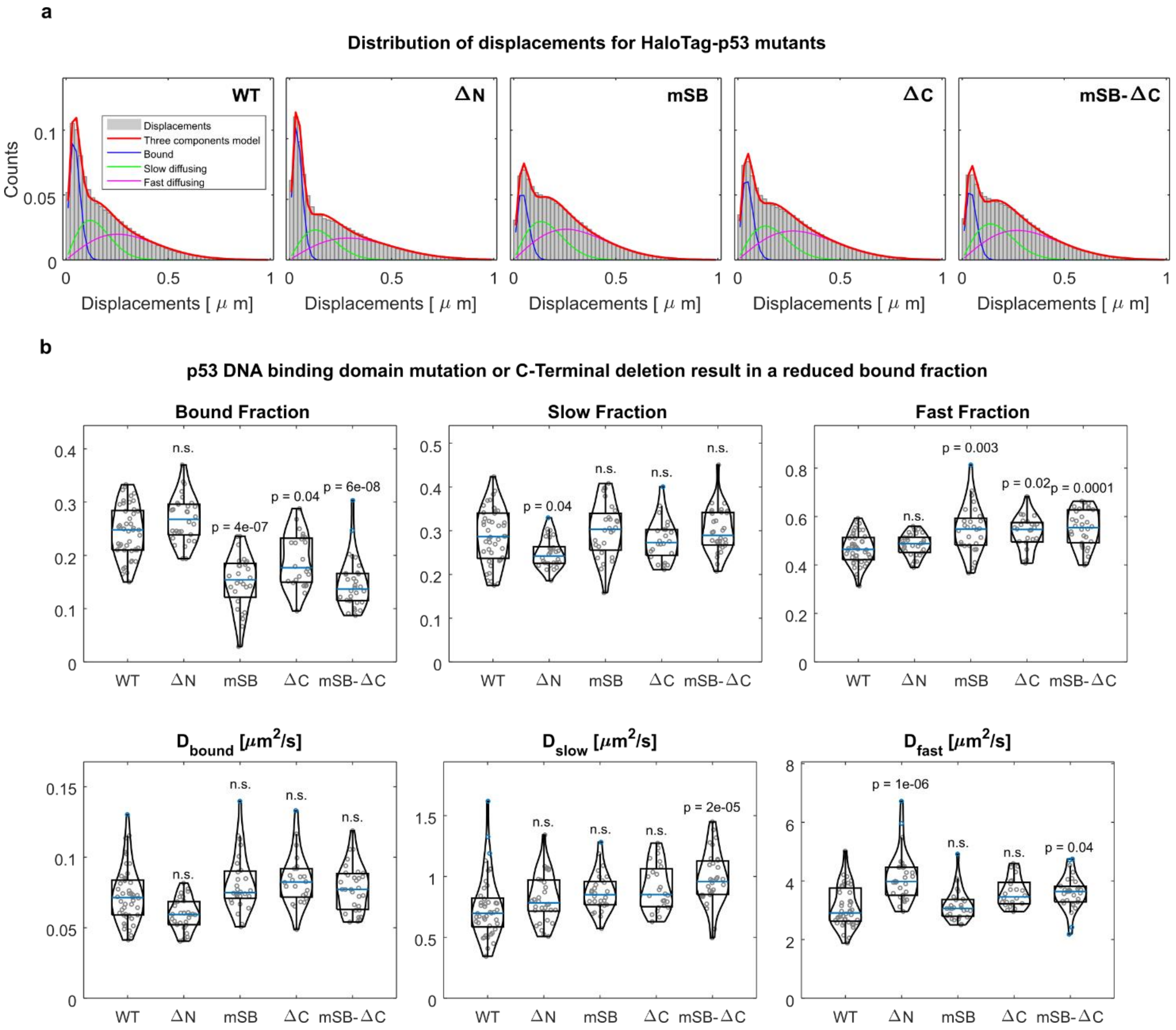


**Supplementary Figure 5.** paSMT analysis of p53 mutants dynamics. (**a**) Distribution of displacements for p53 WT and mutants. (**b**) Single-cell parameters extracted by fitting the distribution of displacements (same data as in Figure 5B, statistical test Kruskal-Wallis, with Bonferroni correction for multiple testing).

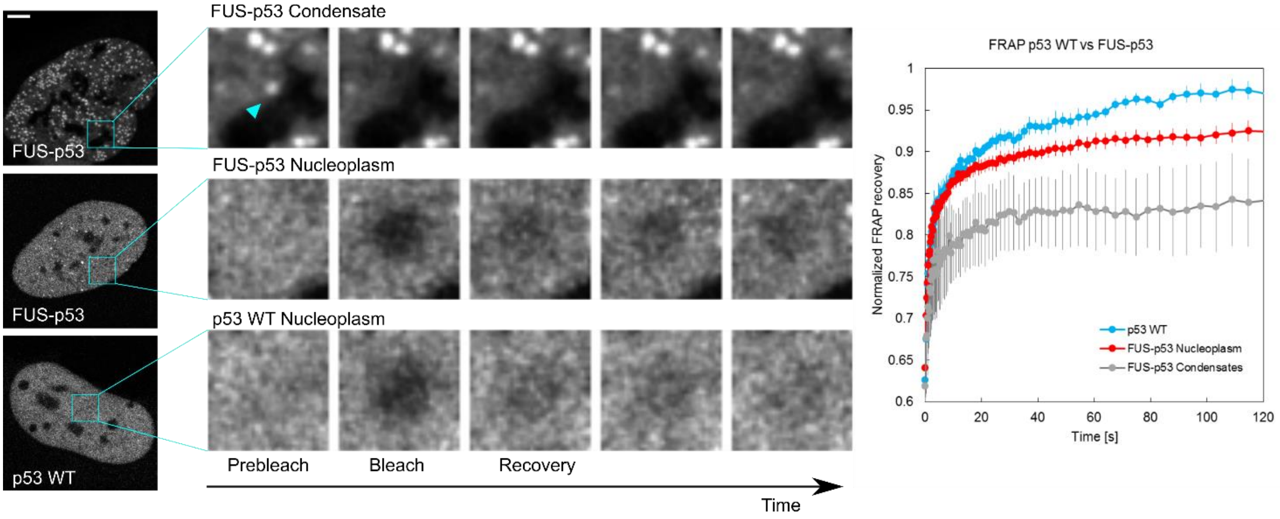


**Supplementary Figure 6. FRAP analysis of p53 WT and FUS-p53 dynamics.** FUS-p53 fluorescence recovery is slowed down compared to p53 WT, both in condensates and outside of them (errorbar SEM, $N_{cells}$= 9, 29, 5 for p53-WT, FUS-p53 nucleoplasm, FUS-p53 condensates respectively).

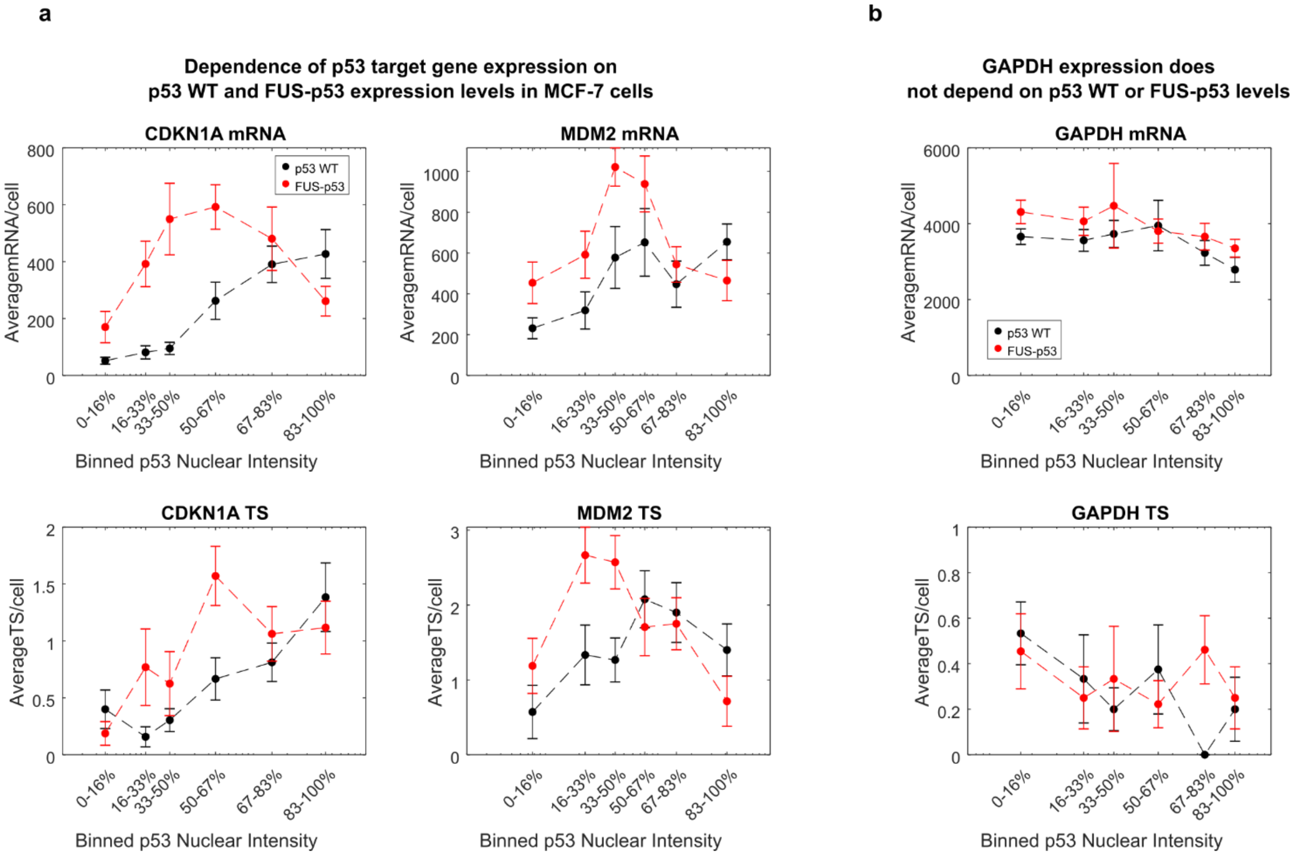


**Supplementary Figure 7. Additional analysis on target gene expression dependence on FUS-p53 levels by smFISH.** (**a**) Simultaneous imaging of HaloTag-p53 nuclear levels and mRNA expression by smFISH highlights that p53 target genes are activated in a bimodal fashion by FUS-p53 also in breast cancer MCF7 cells. (**b**) The expression of the housekeeping gene GAPDH does not depend on p53-wt or FUS-p53 nuclear levels.

### Supplementary Movies Legends.

**Supplementary movie 1.** Exemplary SMT/mSIM acquisition for Halotag, p65, p53, CTCF and H2B. Movies collected at 100fps and displayed at 30fps.

**Supplementary movie 2.** Exemplary paSMT acquisition for Halotag, p65, p53, CTCF and H2B. Movies collected at 100fps and displayed at 30fps.

### Supplementary Note 1: Reconstruction of mSIM images

As described by York et al. (York et al., 2012), in order to reconstruct a super-resolved and optically sectioned image from the 224 individual frames acquired in our mSIM set-up, it is necessary to: (i) identify the position of the illumination spots in the image plane at each acquisition frame; (ii) perform digital pinholing of the individual frames to get rid of out-of focus blur; (iii) fuse together the images through pixel-reassignment that result in a $\sqrt{2}$ increase in the lateral resolution of the microscope. All these steps are performed with custom-written routines in Matlab.

Identification of the illumination spot position: The position at which illumination spots appear in the image, does not necessarily correspond to the points in which the DMD spots illuminate the sample, but rather they are the convolution product of the illumination pattern projected on the sample for the actual distribution of fluorescent labels in the sample. On average, however, the distance between the recorded spots should reflect the average distance between illumination points. Similar to what was performed in York et al., we use this principle to find the lattice vectors that describe the 2D displacement between any two illumination spots. The position of all illumination points in the acquisition stack can be defined by two set of vectors (that need to be found for each individual acquisition): (i) the lattice vectors that describe the displacement between an illumination spot and the two nearest neighbor ones; (ii) the offset vectors, that specify the absolute position of the illumination spot closer to the top-left corner of the image in each of the images of the raw acquisition.

Identification of the lattice vectors. To identify the lattice vector we first localize the positions at which illumination spots appear in the sample by using the ThunderStorm plug-in in ImageJ/FIJI. These coordinates are then used to generate a stack of binary images with ones at the pixels where the illumination spots have been localized and zero elsewhere. The images are then Fourier transformed and the Fourier magnitude images are then averaged together, giving rise to a periodic lattice of peaks, spaced by the inverse of the average spacing between peaks in the real images. Next, we search for peaks in the Fourier dimension, verify that we can find harmonics of the peaks found at lowest spatial frequency, and identify three candidate peaks with the lowest spatial frequency (that shows harmonics). We next verify that the vector sum and differences of the position vector of these identified peaks, also point to a detectable peak. If two vectors satisfy these conditions they are chosen as the lattice vectors in the Fourier space, and are then Fourier transformed to obtain real lattice vectors.

Identification of the offset vectors. Once we have the lattice vectors we can find the position of any other illumination spot in one of the images of the raw stack by knowing the position of one of them (i.e. by finding the position of the top left illumination spot in each them image, the offset vectors). To this scope we translate the expected pattern of illumination over the x and y positions and maximize the autocorrelation between the expected illumination distribution and the experimental one.

Virtual Pinholing and pixel reassignment. To suppress out of focus blur, we next perform virtual pinholing, around the positions of illumination spots detected above. To this scope, we scale the raw images of a factor 10 in both dimensions, and we apply a gaussian mask of 130nm in standard deviation around the position of each detected illumination spot. Next, for every frame in the stack and for every spot position in the image the raw data from a squared region around the illumination spot are copied to the final image matrix in a squared region centered to the original coordinates multiplied by two. This procedure is analogous to move the information from each pixel to half of its distance from the illumination center, the principle on which Image Scanning Microscopy (ISM) is based (York et al., 2012). Summing the result of this procedure for each illumination spot generates the final super-resolved image that is then scaled down by a factor 10 and deconvolved using the Richardson-Lucy algorithm.

### Supplemental Note 2: Correction of the distribution of displacements model for molecules going out of focus.

In SMT only molecules positioned within a slice of thickness of approximately $1\mu\text{m}$ around the focal plane can be localized and tracked. As a consequence, molecules diffusing with faster diffusion coefficients are more likely to diffuse out of focus, resulting in an underestimation of the fraction of molecules involved in this fast diffusion.

As described in the Methods section of the main text, we fit the distribution of displacements with a multi-component diffusion model, described by:

$$p(r)\Delta r = r\Delta r \sum_{i=1}^{3} \frac{f_i}{2D_i\Delta t} \exp\left(-\frac{r^2}{4D_i\Delta t}\right) p(\Delta z, D_i)$$

(Eq S.1)

Here $p(\Delta z, D_i)$, accounts for the probability of molecules of still staying in the slice with limits $-\Delta z$

to $+\Delta z$ around the focal plane. Considering that within this slice a single molecule can have any starting position $z_s$, $p(\Delta z, D_i)$ can be calculated as:

$$p(\Delta z, D) = \frac{1}{2\Delta z}\frac{1}{\sqrt{\pi}}\frac{1}{\sqrt{4Dt}}\int_{-\Delta z}^{+\Delta z}\int_{-\Delta z}^{+\Delta z} e^{-\frac{(z-z_s)^2}{4Dt}}\,\mathrm{d}z\,\mathrm{d}z_s$$

(Eq S.2)

By integrating over $z$, performing the substitution of variables $\theta = \frac{z-z_s}{\sqrt{4Dt}} \rightarrow \mathrm{d}\theta = \frac{\mathrm{d}z}{\sqrt{4Dt}}$, we obtain:

$$p(\Delta z, D) = \frac{1}{2\Delta z}\frac{1}{\sqrt{\pi}}\int_{-\Delta z}^{+\Delta z}\int_{\frac{-\Delta z - z_s}{\sqrt{4Dt}}}^{\frac{+\Delta z - z_s}{\sqrt{4Dt}}} e^{-\theta^2}\,\mathrm{d}\theta\,\mathrm{d}z_s = \frac{1}{2\Delta z}\int_{-\Delta z}^{+\Delta z} \mathrm{erf}\left(\frac{+\Delta z - z_s}{\sqrt{4Dt}}\right) + \mathrm{erf}\left(\frac{+\Delta z + z_s}{\sqrt{4Dt}}\right)\mathrm{d}z_s$$

Where we used the symmetric function $\mathrm{erf}(\theta) = 2/\pi \int_0^{\theta} e^{-\theta^2}\mathrm{d}\theta$

To solve this, we split the integral in two:

$$p(\Delta z, D) = \frac{1}{4\Delta z}\left[\int_{-\Delta z}^{+\Delta z} \mathrm{erf}\left(\frac{+\Delta z - z_s}{\sqrt{4Dt}}\right)\mathrm{d}z_s + \int_{-\Delta z}^{+\Delta z} \mathrm{erf}\left(\frac{+\Delta z + z_s}{\sqrt{4Dt}}\right)\mathrm{d}z_s\right]$$

In the first integral $I$ we can substitute : $\theta = \frac{+\Delta z - z_s}{\sqrt{4Dt}} \rightarrow d\theta = -\frac{dz_s}{\sqrt{4Dt}}$ to yield:

$$I = -\sqrt{4Dt}\int_{\frac{\Delta z}{\sqrt{Dt}}}^{0} \mathrm{erf}(\theta)\,\mathrm{d}\theta = -\sqrt{4Dt}\left[\theta\,\mathrm{erf}(\theta) + \frac{e^{-\theta^2}}{\sqrt{\pi}}\right]_{\frac{\Delta z}{\sqrt{Dt}}}^{0} = 2\Delta z\,\mathrm{erf}\left(\frac{\Delta z}{\sqrt{Dt}}\right) + \sqrt{\frac{4Dt}{\pi}}e^{-\frac{(\Delta z)^2}{Dt}} - \sqrt{\frac{4Dt}{\pi}}$$

Similar the second integral , we substitute: $\theta = \frac{+\Delta z + z_S}{\sqrt{4Dt}} \rightarrow d\theta = +\frac{dz_S}{\sqrt{4Dt}}$

$$II = \sqrt{4Dt}\int_{0}^{\frac{\Delta z}{\sqrt{Dt}}} \mathrm{erf}(\theta)\,\mathrm{d}\theta = 2\Delta z\,\mathrm{erf}\left(\frac{\Delta z}{\sqrt{Dt}}\right) + \sqrt{\frac{4Dt}{\pi}}e^{-\frac{(\Delta z)^2}{Dt}} - \sqrt{\frac{4Dt}{\pi}}$$

Summing up the terms for $I$ and $II$, we obtain:

$$p(\Delta z, D) = \frac{1}{2\Delta z}\left(2\Delta z\,\mathrm{erf}\left(\frac{\Delta z}{\sqrt{Dt}}\right) + \sqrt{\frac{4Dt}{\pi}}e^{-\frac{(\Delta z)^2}{Dt}} - \sqrt{\frac{4Dt}{\pi}}\right)$$

Which is the final expression for $p(\Delta z, D)$ used in Eq. S1.

### Supplemental Tables.

| smRNA FISH probes for ***CDKN1A*** (*HuluFISH,* PixelBiotech GmbH) | |
|---|---|
| Name | Sequence |
| CDKN1A_211 | CTGCCGCAGAAACACCTGT |
| CDKN1A_0 | CCCAGCCGGTTCTGACAT |
| CDKN1A_18 | GCATGGGTTCTGACGGACATC |
| CDKN1A_85 | ATCACAGTCGCGGCTCAGCT |
| CDKN1A_160 | TCCAGTGGTGTCTCGGTGA |
| CDKN1A_219 | CGTGGGAAGGTAGAGCTTGGG |
| CDKN1A_255 | TCCTCCCAACTCATCCCG |
| CDKN1A_305 | TCTTCCTCTGCTGTCCCCTGC |
| CDKN1A_345 | GCGAGGCACAAGGGTACAAGA |
| CDKN1A_366 | TTCAGCCTGCTCCCCTGA |
| CDKN1A_393 | TGAGAGTCTCCAGGTCCACC |
| CDKN1A_427 | TCTGTCATGCTGGTCTGCC |
| CDKN1A_447 | CCGGCGTTTGGAGTGGTAGAA |
| CDKN1A_11 | TTCCAGGACTGCAGGCTTCC |
| CDKN1A_41 | ATGTAGAGCGGGCCTTTGAG |
| CDKN1A_121 | GCCAGGGTATGTACATGAGGA |
| CDKN1A_187 | CTCTCATTCAACCGCCTAGTT |
| CDKN1A_208 | TGCCCAGCACTCTTAGGAAC |
| CDKN1A_251 | ACACGGGATGAGGAGGCTTTA |
| CDKN1A_313 | GGAGGAGGAAGTAGCTGGCAT |
| CDKN1A_335 | TACCACCCAGCGGACAAGTG |
| CDKN1A_367 | AGCGATGGGAAGGAGCCACAC |
| CDKN1A_393 | GGGTGAATTTCATAACCGCCT |
| CDKN1A_414 | GGTCTGAGTGTCCAGGAAAGG |
| CDKN1A_460 | CCCTTCAAAGTGCCATCTGTT |
| CDKN1A_483 | ATGATGCCCCCACTCGGTGAG |
| CDKN1A_530 | CACCCTGCCCAACCTTAGAG |
| CDKN1A_550 | GCTGTGCTCACTTCAGGGT |
| CDKN1A_576 | TACCAGGTCCCCAGCTCA |
| CDKN1A_594 | GGGTATCAAGAGCCAGGAGGG |
| CDKN1A_620 | CCCCTGCCTTCACAAGACA |
| CDKN1A_676 | TGCAGGTCAGAGGGGCCATGA |
| **smRNA FISH probes for *GAPDH*** (*HuluFISH,* PixelBiotech GmbH) | |
| GAPDH_35 | CGGCTGGCGACGCAAAAGAAG |
| GAPDH_57 | TGGTGTCTGAGCGATGTGGC |
| GAPDH_77 | CGACCTTCACCTTCCCCA |
| GAPDH_102 | GCCCAATACGACCAAATCCG |
| GAPDH_132 | CCAGAGTTAAAAGCAGCCCTG |
| GAPDH_235 | CGGTGCCATGGAATTTGCC |
| GAPDH_254 | CTTCCCGTTCTCAGCCTTGA |
| GAPDH_291 | TCCTGGAAGATGGTGATGGGA |

| | |
|---|---|
| GAPDH_317 | CCCACTTGATTTTGGAGGGAT |
| GAPDH_345 | TCCACGACGTACTCAGCGCCA |
| GAPDH_384 | CCAGCCTTCTCCATGGTG |
| GAPDH_424 | GGCAGAGATGATGACCCTTTT |
| GAPDH_452 | TGACGAACATGGGGGCATCAG |
| GAPDH_499 | GCTGATGATCTTGAGGCTGTT |
| GAPDH_530 | TGCTAAGCAGTTGGTGGTGC |
| GAPDH_575 | GTCCTTCCACGATACCAAAGT |
| GAPDH_601 | GTGATGGCATGGACTGTGGT |
| GAPDH_629 | GGGCCATCCACAGTCTTCT |
| GAPDH_648 | TCACGCCACAGTTTCCCGGAG |
| GAPDH_674 | TGATGTTCTGGAGAGCCCCGC |
| GAPDH_739 | CTTCCCGTTCAGCTCAGG |
| GAPDH_786 | CCACCACTGACACGTTGGCA |
| GAPDH_811 | AGGTTTTTCTAGACGGCAGGT |
| GAPDH_841 | CACCACCTTCTTGATGTCATC |
| GAPDH_888 | TCAGTGTAGCCCAGGATGCCC |
| GAPDH_925 | GGTGTCGCTGTTGAAGTCAGA |
| GAPDH_947 | CAGCGTCAAAGGTGGAGGAGT |
| GAPDH_978 | ACAAAGTGGTCGTTGAGGGCA |
| GAPDH_1018 | GCTGTAGCCAAATTCGTTGTC |
| GAPDH_1061 | ACTCCTTGGAGGCCATGTGGG |
| GAPDH_1107 | TCTCTTCCTCTTGTGCTCTTG |
| GAPDH_1145 | ACTGAGTGTGGCAGGGACTCC |
| **smRNA FISH probes for *MDM2*** (*Design Ready Stellaris,* Biosearch Technologies) | |
| MDM1 | TGTTGGTATTGCACATTTGC |
| MDM2 | TACAGCACCATCAGTAGGTA |
| MDM3 | CGAAGCTGGAATCTGTGAGG |
| MDM4 | TAAGTGTCTTTTTGTGCACC |
| MDM5 | TAGTCATAATATACTGGCCA |
| MDM6 | ATGTTGTTGCTTCTCATCAT |
| MDM7 | TGGCACGCCAAACAAATCTC |
| MDM8 | CTGTGCTCTTTCACAGAGAA |
| MDM9 | TTCCTGTAGATCATGGTATA |
| MDM10 | CTGCTGATTGACTACTACCA |
| MDM11 | TGATCACTCCCACCTTCAAG |
| MDM12 | GGTAGATGGTCTAGAAACCA |
| MDM13 | CTAATTGCTCTCCTTCTAGA |
| MDM14 | GTCGTTCACCAGATAATTCA |
| MDM15 | CTATCAGATTTGTGGCGTTT |
| MDM16 | TATTACACACAGAGCCAGGC |
| MDM17 | TTCTTTCACAACATATCTCC |
| MDM18 | CCTGTAGATTCACTGCTACT |

| | |
|---|---|
| MDM19 | CACTTACACCAGCATCAAGA |
| MDM20 | TCCAACCAATCACCTGAATG |
| MDM21 | CTGATCTGAAACTGAATCCT |
| MDM22 | CCTTCTTCACTAAGGCTATA |
| MDM23 | CTGCCTGATACACAGTAACT |
| MDM24 | ATTGCATGAAGTGCATTTCC |
| MDM25 | AATGTGATGGAAGGGGGGGA |
| MDM26 | GTGTTGAGTTTTCCAGTTTG |
| MDM27 | ACTCTCTGGAATCATTCACT |
| MDM28 | TCATCATTTTCCTCAACACA |
| MDM29 | TGTGATTGTGAAGCTTGTGT |
| MDM30 | GGCTGAGAATAGTCTTCACT |
| MDM31 | ACTCTTTCACATCTTCTTGG |
| MDM32 | AAGGGGCAAACTAGATTCCA |
| MDM33 | TCACACAAGGTTCAATGGCA |
| MDM34 | TAAGATGTCCTGTTTTGCCA |
| MDM35 | TTTGCACATGTAAAGCAGGC |
| MDM36 | TTGGTTGTCTACATACTGGG |
| MDM37 | AGGGGAAATAAGTTAGCACA |
| MDM38 | ATTCTCTTATAGACAGGTCA |

**Supplementary Table 1. List of Fluorescently labeled oligos used for smFISH**
